# The same plasticity rule rescues networks built from one conductance set and destabilises networks built from another

**DOI:** 10.64898/2026.09.15.751713

**Authors:** Filippo Groppi, Jessica Ferrari

## Abstract

Whether synaptic plasticity stabilises or destabilises a recurrent circuit is usually treated as a question about the rule. We find that in a conductance-based model it is not answered by the rule alone.

The model is a simulated bursting culture: 48 excitatory and 12 inhibitory stomatogastric-ganglion neurons, sparsely and recurrently connected. The recurrent synapses carry pair-based spike-timing-dependent plasticity under a fixed homeostatic budget on each cell’s total incoming excitatory conductance. We ran one net-depressing rule, at one inhibition level, on unselected random wirings, with a survival criterion fixed before the runs.

Two excitatory populations built from different published conductance sets gave **opposite outcomes**. In one, the network collapsed without the rule in 33 of 36 wirings and the rule rescued **31 of those 33**. In the other, the network survived without the rule in 11 of 12 wirings and the rule **eliminated 6 of those 11**. The only model parameter changed between the two is the excitatory conductance set, and that change also alters their intrinsic dynamics.

We then asked what carries the reversal. Across seven conductance sets, elimination increases with a cell’s own firing rate, and cell identity does not predict it — a follower-type cell firing at pacemaker rates behaves like a pacemaker. But moving the firing rate inside a single set by injected current reverses the same quantity, which rules out firing rate as a sufficient explanation.

Substituting one conductance at a time from the rescued set into the other, over the six channels that differ, no single substitution transferred the rescue (2 of 42 pooled) while substituting all eight did so in 6 of 6 (Fisher exact p = 2.3 × 10^−6^). **The sign reversal cannot be attributed to any single tested conductance, and firing rate alone cannot account for it**.

All results are computational and concern one circuit model, one plasticity rule and one homeostatic constraint. Predictions and falsifiers were committed to a version-controlled record before the corresponding runs, with three departures from that order recorded in §9.

## 1. Introduction

A large literature asks whether activity-dependent synaptic plasticity stabilises recurrent circuits or drives them to pathology, and answers are usually framed as properties of the rule alone: additive spike-timing-dependent plasticity with hard bounds is unstable and multiplicative forms are stable (Song, Miller & Abbott, 2000; van Rossum, Bi & Turrigiano, 2000; Morrison, Diesmann & Gerstner, 2008), and homeostatic terms restore stability provided they act fast enough — a constraint Zenke & Gerstner (2017) show is severe, because measured synaptic scaling is orders of magnitude slower than the Hebbian instability it must contain. The implicit unit of analysis is the rule, sometimes the network, rarely the neuron.

The homeostatic constraint used here is an idealisation of synaptic scaling (Turrigiano et al., 1998; Turrigiano, 2008): the total incoming excitatory conductance per cell is held to a fixed budget, instantaneously rather than over hours. That makes the budget a strong and biologically fast version of scaling, which is the regime in which the rule’s effect is most constrained and, as it turns out, most cell-dependent.

If stability is not a property of the rule alone, testing that requires a model whose intrinsic conductances are catalogued and separately manipulable. The stomatogastric tradition provides one.

Model neurons in that tradition make a different unit available. Prinz, Bucher & Marder (2004) catalogued conductance sets that produce pyloric-like activity, and the same catalogue contains cells whose intrinsic firing rates differ by more than an order of magnitude at the same synaptic drive. If the outcome of a plasticity rule depends on the cell it runs on, this catalogue is the place it would show.

We ran one rule — pair-based, net-depressing, under a fixed homeostatic budget on total incoming excitatory conductance — on populations built from seven of those conductance sets, at one inhibition level, on unselected random wirings, with a survival criterion fixed in advance. The question was not whether the rule is stabilising. It was whether that question has a single answer inside one circuit. It does not: in this model **the effect of a plasticity rule is not determined by the rule alone, but depends on intrinsic cellular properties**.

Two features of the design are worth stating before the results. First, every prediction, falsifier and stopping rule was committed to a version-controlled record before the corresponding run, and the index in §9 gives the registration commit and the report commit for each result; §8 lists every prediction that failed, including all of ours. Second, the quantities are conditional probabilities on a 2 × 2 table, and the two conditionals have denominators that trade off against each other: the fraction of dying networks that the rule rescues can only be measured where the control dies, and the fraction of surviving networks that it eliminates only where the control lives. This is not merely a statistical complication: it follows directly from the 2 × 2 structure of the outcome, and it governs which cell can answer which question.

## 2. Results

### 2.1 The rule redistributes conductance rather than removing it

Under the homeostatic budget the rule does not reduce total recurrent excitation; it concentrates it. On cells driven above rheobase, where firing does not depend on the recurrent connections at all, between a fifth and a quarter of the synapses are driven to zero — **22% on the cortical model** (per-wiring range 0.18–0.26) and **15% on the Hodgkin–Huxley model** (0.14–0.17) — while the population total falls **2.15%** and **10.15%** below its budget respectively, and **no network dies on either arm: 0 of 4 wirings on each of the two cell models**. The much smaller deficits recorded elsewhere in this programme (−1.4% and −0.6%) belong to the STG cells of a different row and are not these.

That is the boundary case that locates the effects below: where activity does not need the recurrence, redistributing it changes nothing we measured.

**Figure 1.**
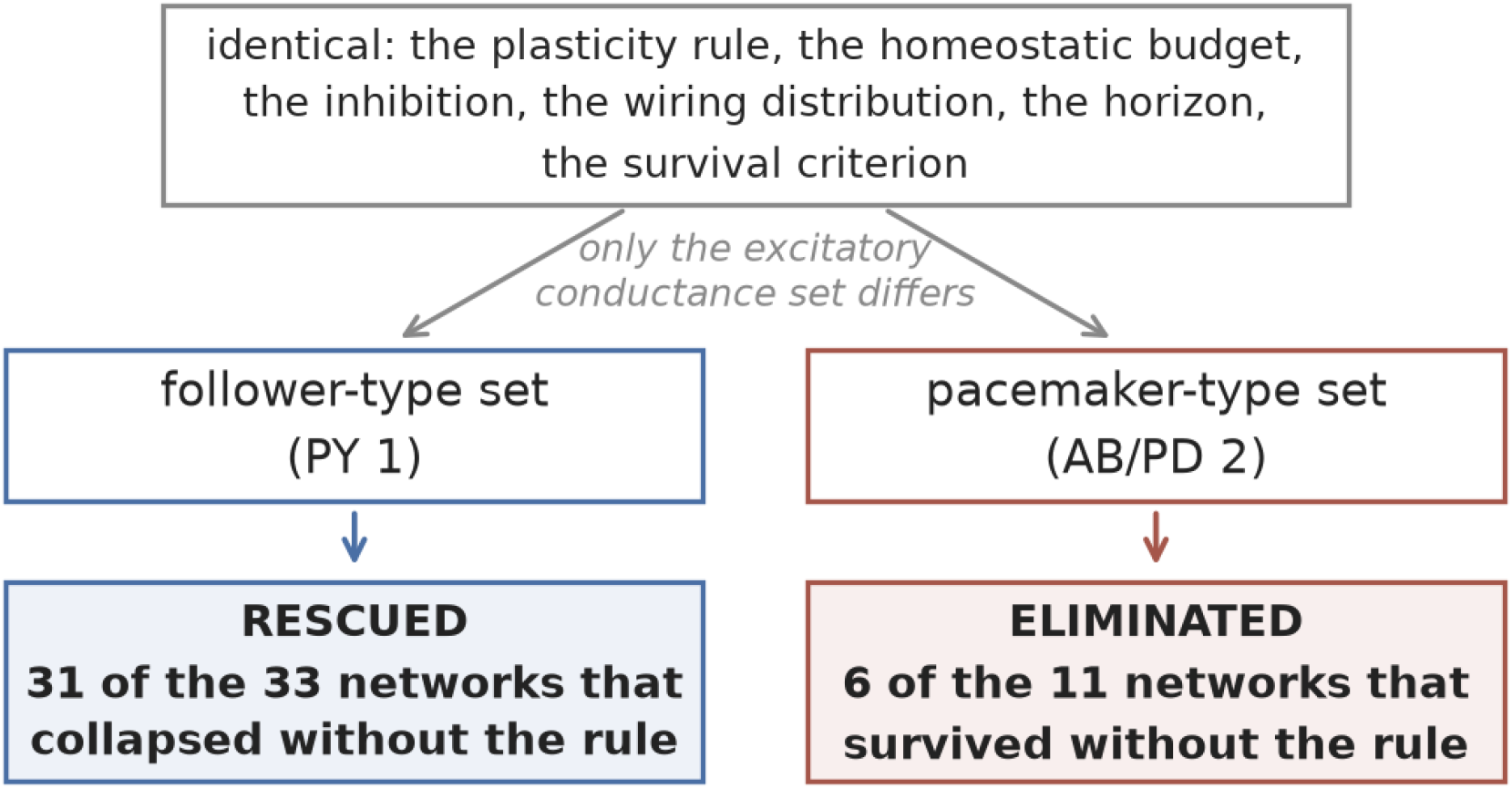
The design and its result in one view. Everything above the split is identical between the two populations — the plasticity rule, the homeostatic budget, the inhibitory conductance, the wiring distribution, the simulation horizon and the survival criterion — and the only model parameter that differs is the excitatory conductance set, which also changes the populations’ intrinsic dynamics. The outcome inverts: the rule rescues most of the networks that collapse without it in one population and eliminates the majority of those that survive without it in the other. Numbers are read from the recorded runs; source values in `figures/fig1_concept.csv`.

**Figure 2.**
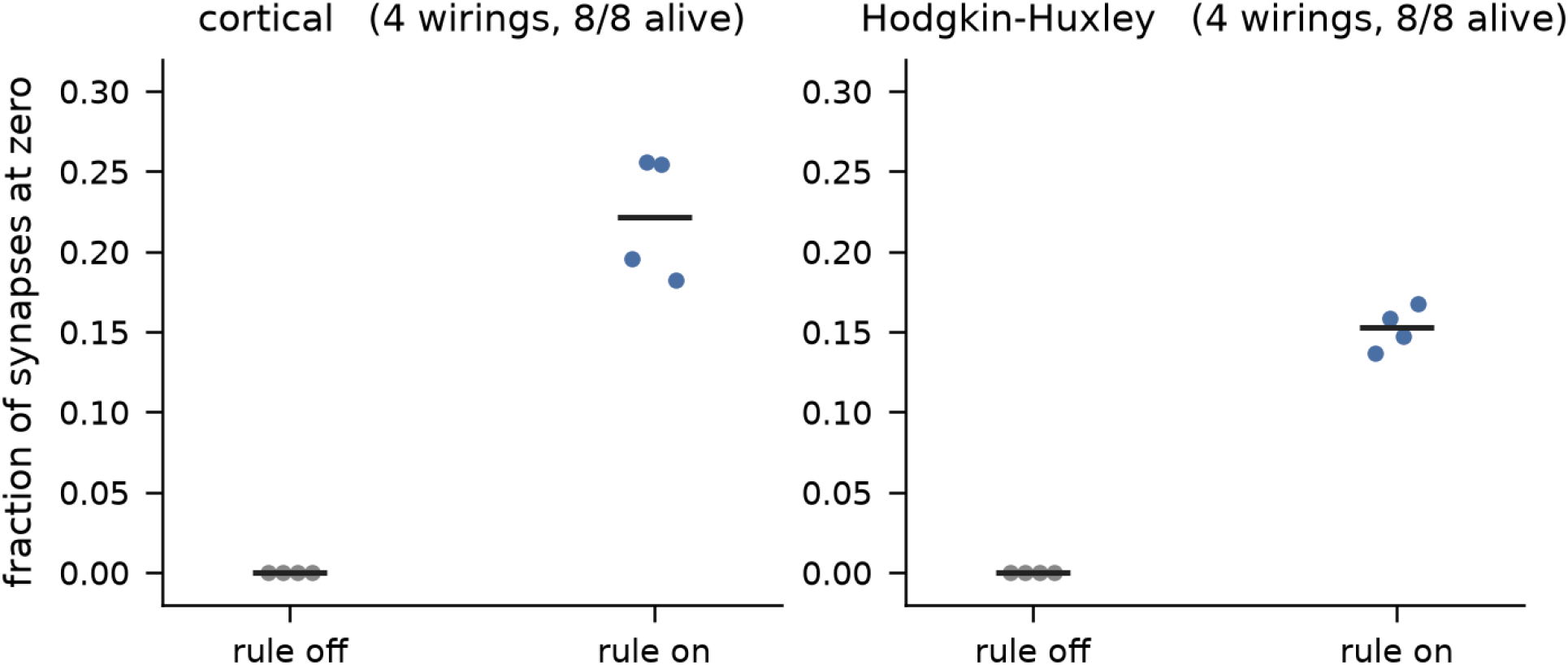
Fraction of recurrent excitatory synapses driven to zero after 60 s, with the rule off and with it on, for two cell models driven above rheobase (four wirings each; points are wirings, bars are arm means). The total incoming conductance is held to a fixed budget throughout, so the movement to zero is redistribution and not removal. All eight networks survive in both arms. Source values in `figures/fig2_redistribution.csv`.

### 2.2 The same rule has opposite effects on two conductance sets

Two excitatory populations, identical in every respect except the published conductance set they are built from, at the same inhibitory conductance g_ie = 0.045, on unselected random wirings, with survival defined in advance as at least 24 of 48 excitatory cells firing in the final 10 s of a 35 s run.

**On PY 1**, a follower-type set, the control arm collapses in 33 of 36 wirings, and with the rule running:

|  | rule on, alive | rule on, dead |
| --- | --- | --- |
| rule off, alive | 3 | 0 |
| rule off, dead | 31 | 2 |

**P(alive with the rule │ dead without it) = 31/33 = 0.939**, Wilson 95% CI **[0.804, 0.983], p = 6.5 × 10**^**−8**^. Of the three wirings that would have survived anyway, the rule eliminates none.

**On AB/PD 2**, a pacemaker-type set at the same inhibition, the control arm survives in 11 of 12 wirings and the rule **eliminates 6 of those 11** (0.545, Wilson [0.280, 0.787]) while rescuing the single wiring that died . At g_ie = 0.060 it eliminates 4 of 11; at g_ie = 0.030 nothing dies on either arm and the table is empty, which shows the effect needs something to act on.

The inhibition, the rule, the budget, the wiring distribution, the horizon and the detector are identical between the two. **The only model parameter changed between these two populations is the excitatory conductance set, and that change also changes their intrinsic dynamics** — they differ by more than an order of magnitude in firing rate at the same synaptic drive, which is the subject of §2.3 and the caveat on reading §2.2 as an isolated comparison.

#### Both cells were measured under two detectors

The registered one asks whether a cell fired at any point in the final 10 s; a stricter variant requires the quorum in the final 5 s alone. **On both cells, at 35 s, the two detectors agree on every wiring**.

**Figure 3.**
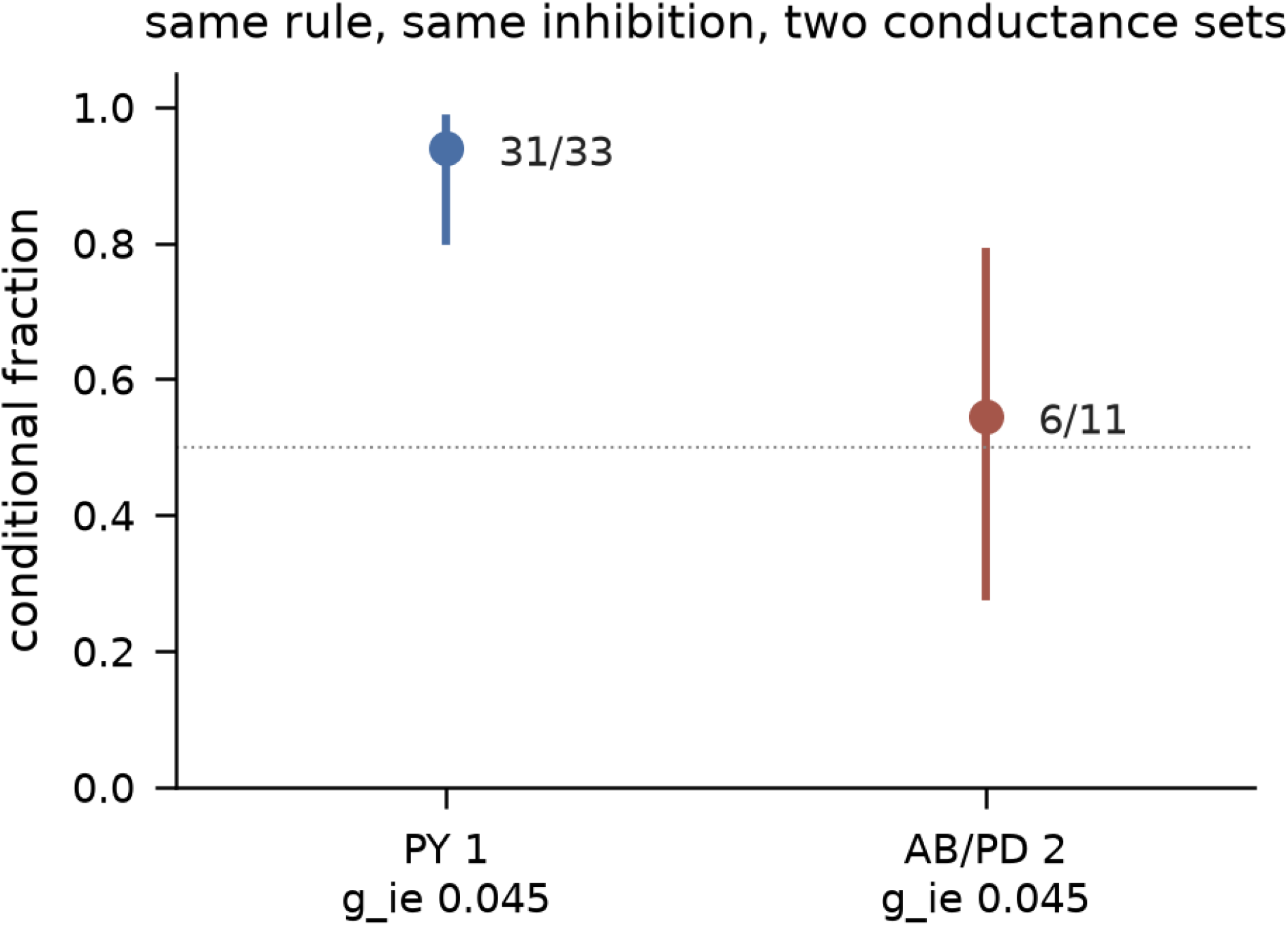
The two conditional outcomes with Wilson 95% intervals: for PY 1, the fraction of networks that collapsed without the rule and survived with it; for AB/PD 2, the fraction that survived without the rule and were eliminated by it. Same inhibition, same rule, same wiring distribution; the conductance set is the only difference. The dotted line marks 0.5. Source values, including the g_ie 0.030 and 0.060 tables, in `figures/fig3_two_signs.csv.`

### 2.3 Elimination increases with firing rate across the sampled conductance sets

Seven conductance sets at g_ie = 0.045, each contributing its own control-arm firing rate measured in the same runs, all at a 35 s horizon. The comparable statistic across cells is the fraction of surviving wirings the rule eliminates: unlike (rescued − eliminated), it has no ceiling set by the control’s mortality, which is itself the ordering variable.

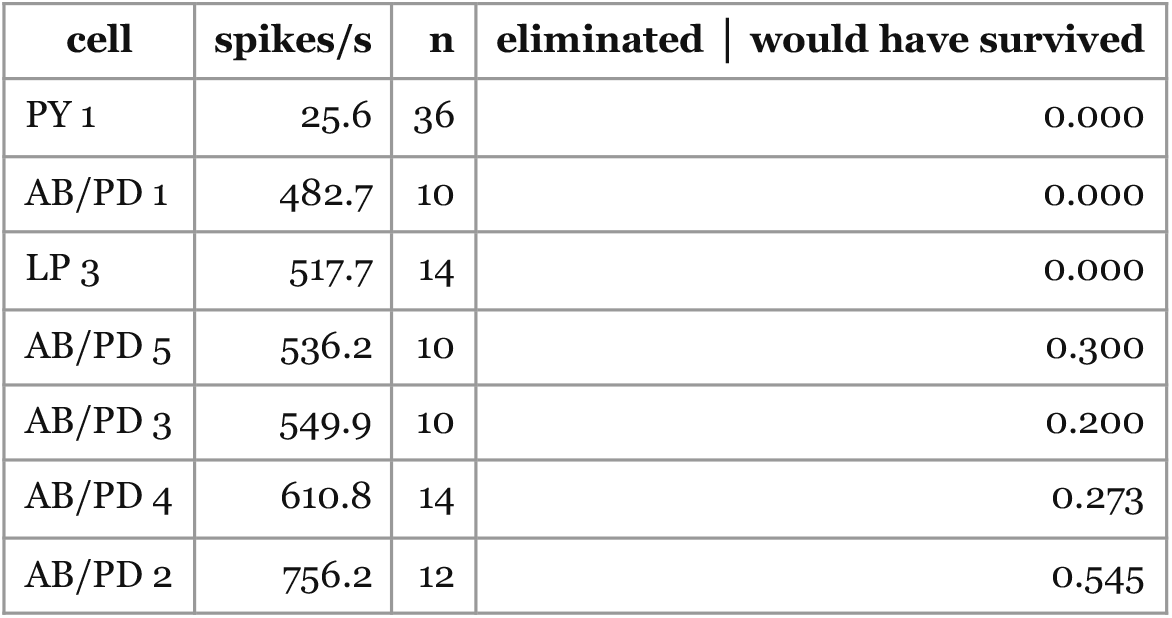

#### Spearman +0.852, p = 0.0286

Within the seven sets sampled here, elimination increases monotonically with intrinsic firing rate.

#### Cell identity does not predict it

LP 3 is a follower cell, like PY 1, and was named in the registration as the case that would separate a firing-rate account from an identity account, because it fires at 517.7 spikes/s — next to the pacemakers and nowhere near PY 1’s 25.6. It behaves like the pacemakers at that rate: 0.000 eliminated, identical to AB/PD 1, and nothing like PY 1.

Two limits belong with this number. **The relationship is carried by the low end**: excluding PY 1 leaves Spearman +0.812 at p = 0.072 over six cells, and PY 1 sits at 25.6 spikes/s with the next cell at 482.7, a nineteenfold gap that the sampled catalogue does not fill. And **AB/PD 2’s own elimination excess is not individually significant** — 6 eliminated against 1 rescued is seven discordant wirings, p = 0.125 — so the cell at the high end supports the relationship as a point estimate and not as a test.

**Figure 4.**
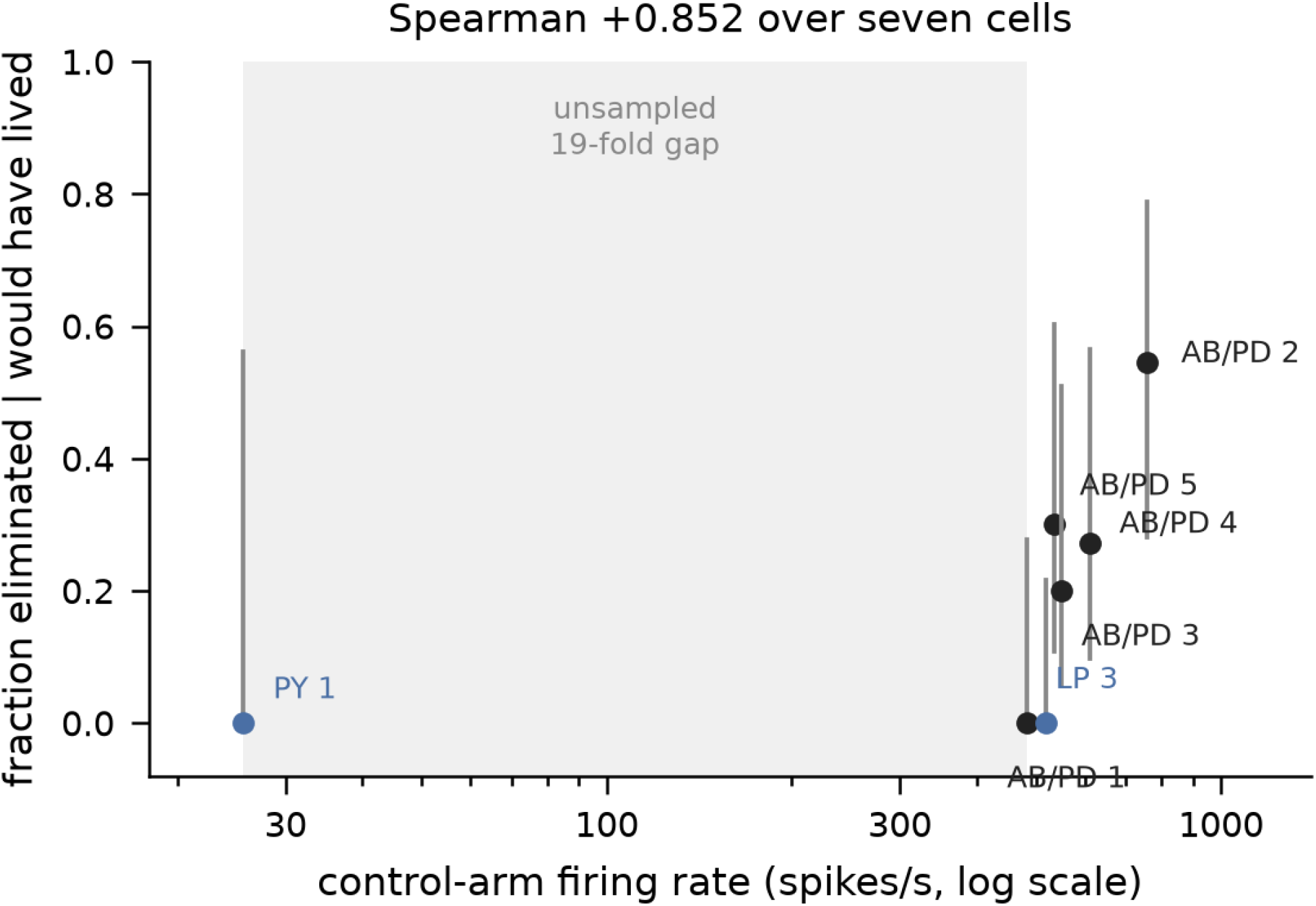
Fraction of surviving networks eliminated by the rule, against the control arm’s own firing rate, for seven conductance sets on a logarithmic axis; bars are exact Wilson 95% intervals and the shaded band is the unsampled interval between the slowest set and the next. The two follower-type sets are labelled in colour: one is the slowest cell and one fires at pacemaker rates. All seven were measured at the same 35 s horizon. Source values in `figures/fig4_ordering.csv.`

### 2.4 Changing the firing rate within one cell reverses the effect

The relationship in §2.3 is across conductance sets, and it does not survive a manipulation inside one. Injecting a constant hyperpolarising current into the excitatory population of AB/PD 1 — the same conductance set, the same wirings, only the drive moved — takes the elimination fraction from **0.000 at its natural rate of 483 spikes/s to 1.000 at the driven setting**, where the rule eliminated all three wirings that would otherwise have survived.

Across cells, elimination rises with rate. Within this cell, it rises as the rate falls. A cross-cell association is not a within-cell causal manipulation, and here the two disagree: **the manipulation falsifies firing rate as a sufficient explanatory variable for the effect**. §2.3 records an association across the sampled sets; it does not license a causal reading in terms of rate.

### 2.5 Fragility and rescuability dissociate

The same manipulation separates two things that §2.3 conflated, because the only rescued cell there is also the only slow one.

| condition | control-arm rate | control dead | rescued |
| --- | --- | --- | --- |
| AB/PD 1, no injected current | ~483 spikes/s | 0/10 | — |
| AB/PD 1, $-0.03 \mu\text{A}/\text{cm}^2$ | — | 9/12 | 0/9 |
| AB/PD 1, $-0.05 \mu\text{A}/\text{cm}^2$ | 132 (first 5 s) | 12/12 | 0/12 |
| PY 1, no injected current | 25.6 spikes/s | 33/36 | **31/33** |

#### Fragility follows the rate within one fixed cell

0 of 10 controls collapse at baseline and 12 of 12 at the driven setting. **Rescuability does not**. At matched fragility the rule rescues PY 1 in 31 of 33 opportunities and the driven cell in **0 of 21** (**p = 4.9 × 10**^**−13**^).

#### Slow firing predicts fragility, but not rescue

##### The state of the driven preparation limits this comparison

The 132 spikes/s above is measured over the first 5 s, before any collapse; over the whole run the driven preparation averages **19 spikes/s and produces zero network bursts**. It is therefore matched to the follower cell on **control mortality and not on dynamical state** — it has left the bursting regime, and the follower has not. What the comparison licenses is that equalising how often the control dies does not equalise rescuability; it does not license the stronger reading that two otherwise equivalent preparations differ only in conductances.

**Figure 5.**
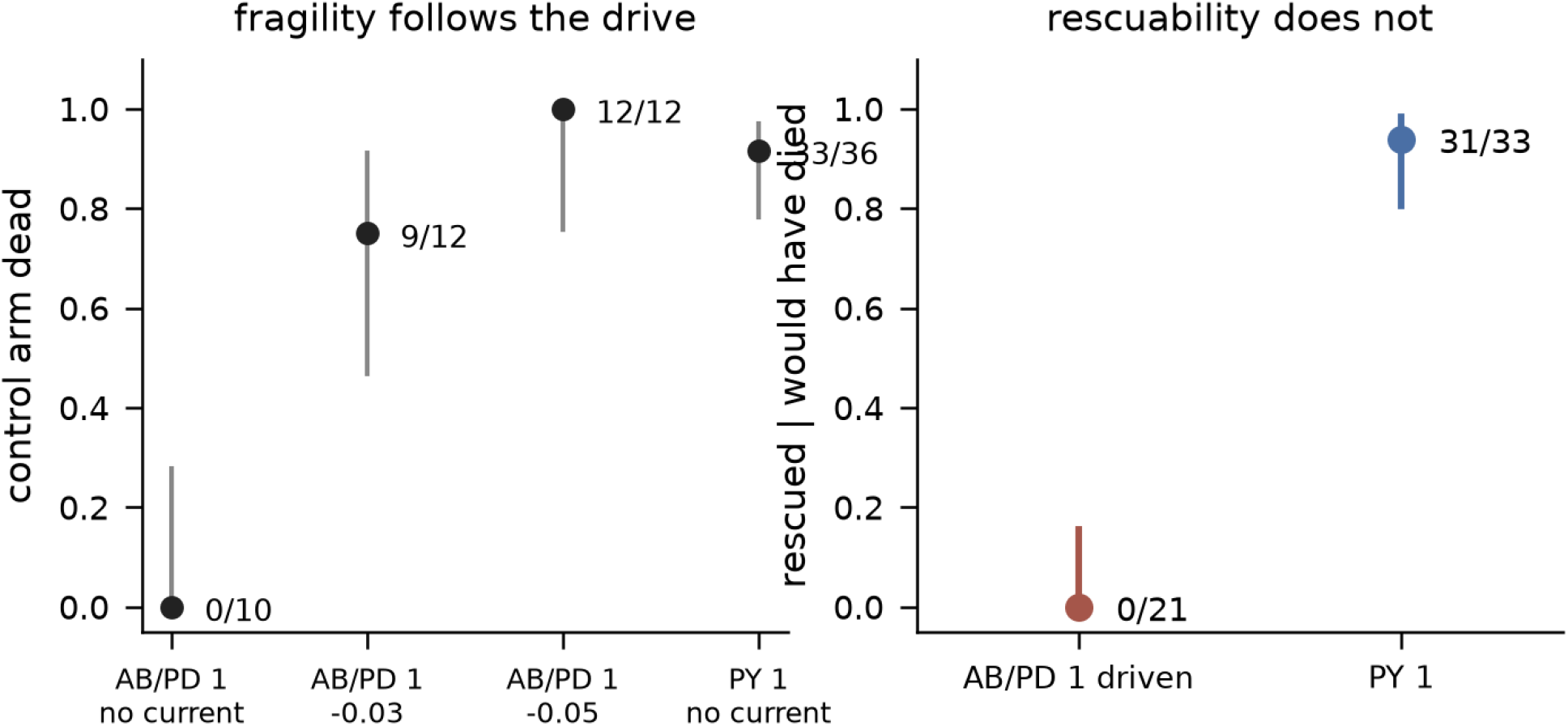
Left: the fraction of control networks that collapse, for AB/PD 1 at its natural drive and under two injected currents, and for PY 1; the drive manipulation moves one conductance set from never collapsing to always collapsing. Right: the fraction of collapsing networks that the rule rescues, for the driven cell and the follower, at matched control mortality. Bars are Wilson 95% intervals. Source values in `figures/fig5_fragility_vs_rescue.csv.`

A control registered after the primary analysis, because the primary analysis was circular: the per-wiring rate first used to relate rate to death was averaged over the whole run, and a wiring that dies has a low average because it died. Re-running the control arm for 5 s — the same deterministic opening, before any death — gives a rate that cannot depend on survival, and the three wirings that survived are exactly the three with the highest early rates (complete rank separation; exact one-sided p = 1/C(12,3) = 0.0045).

### 2.6 No single tested conductance substitution transfers the rescue

We substituted PY 1’s value for AB/PD 1’s, one channel at a time, exhaustively over the eight conductances of the model, at a drive re-chosen per arm so that each arm’s own control dies. Two of the eight conductances are identical in the two cells (I_CaT and I_A), so six substitutions are informative; one of the identical pair is retained as a gate and reproduces the unsubstituted cell bit for bit.

The positive control is the complete substitution, which is PY 1 by construction and is run at the same drive, wirings and horizon as the single substitutions.

| substitution | drive ($\mu\text{A}/\text{cm}^2$) | control dead | rescued | Fisher vs 0/21 |
| --- | --- | --- | --- | --- |
| **all eight (= PY 1)** | −0.03 | 6/7 | **6/6** | **$< 10^{-4}$** |
| I_Na, fast sodium | −0.03 | 7/7 | 0/7 | 1.000 |
| I_CaS, slow calcium | −0.03 | 7/7 | 0/7 | 1.000 |
| I_KCa, calcium-activated potassium | −0.03 | 7/7 | 1/7 | 0.250 |
| I_Kd, delayed-rectifier potassium | −0.03 | 7/7 | 1/7 | 0.250 |
| I_H, hyperpolarisation-activated inward | −0.30 | 7/7 | 0/7 | 1.000 |
| I_leak, leak | −0.14 | 7/7 | 0/7 | 1.000 |

#### Six single substitutions pooled rescue 2 of 42; the complete substitution rescues 6 of 6; p = 2.3 × 10^−6^

The complete substitution’s own control rate is 32 spikes/s, and **two** single substitutions are rate-matched to it: I_Na at 23 and I_KCa at 27 spikes/s, both with 7 of 7 controls dead. Comparing the complete substitution against those two directly gives **p = 0.0006 and p = 0.005**. The other four are not rate-matched (1, 75, 18 and 16 spikes/s) and their comparisons are not offered as matched ones.

#### We do not identify a mechanism, and we state the result as an absence

The decomposition is exhaustive over single substitutions and empty: **no single tested conductance substitution was sufficient to transfer the rescue phenotype**. Whether any is *necessary* is a different question, addressed by the complementary design (substitute all eight, then restore one at a time) and not by this one.

**Figure 6.**
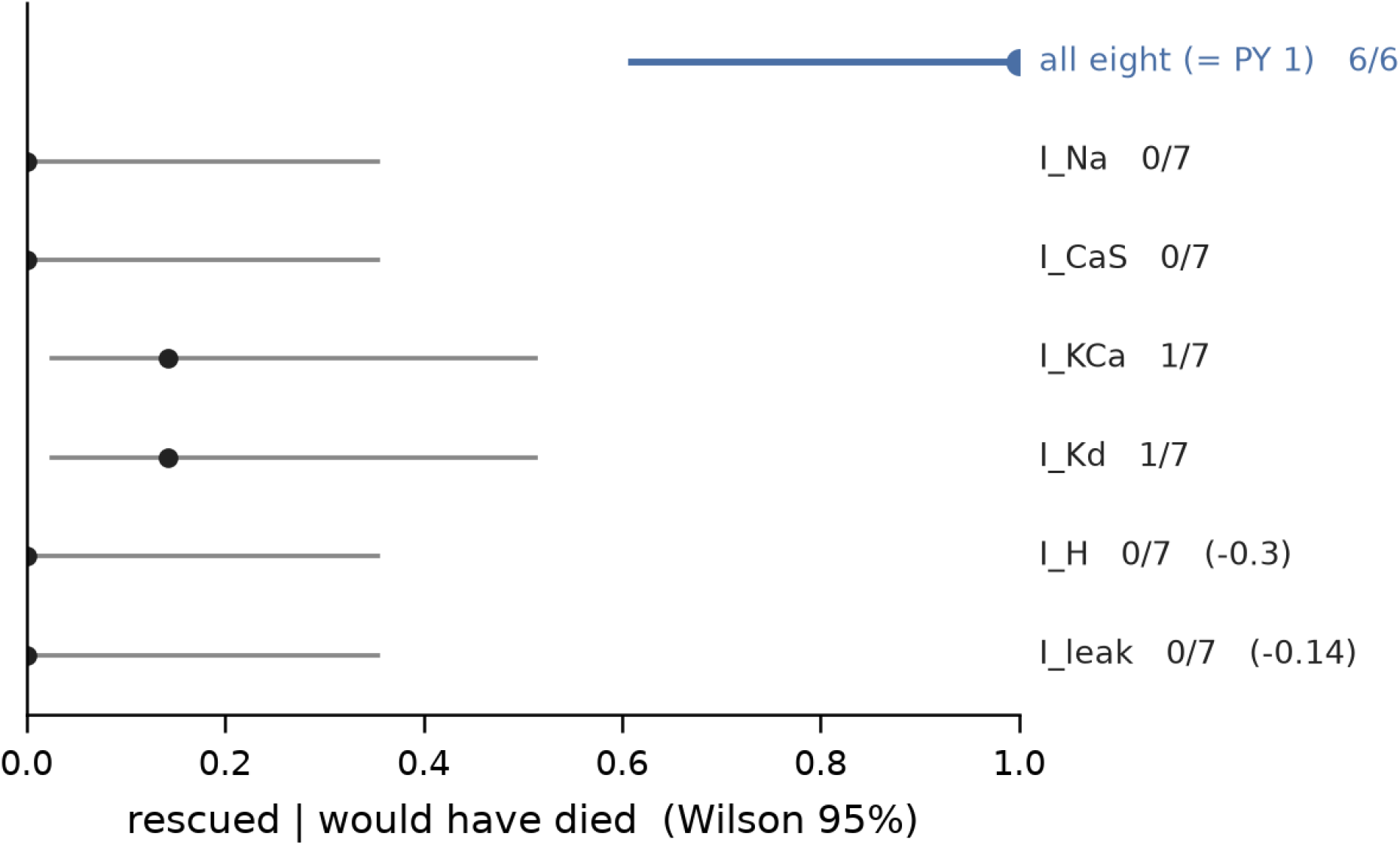
Fraction of collapsing networks rescued by the rule, for the complete eight-channel substitution and for each informative single substitution, with Wilson 95% intervals; seven wirings per arm and each arm at a drive chosen so that its own control collapses, printed where it differs from the common one. The two channels identical in both conductance sets are used as gates and are not plotted. Source values in `figures/fig6_decomposition.csv.`

## 3. Discussion

*In short: within one simulated circuit, the same plasticity rule has opposite effects on two conductance sets; firing rate orders the destructive direction across the sampled sets but is falsified as a sufficient explanation within one; and no single conductance substitution transfers the protective one. We identify no mechanism, and make no claim beyond this model*.

In this model the outcome of a plasticity rule is not determined by the rule alone, but depends on intrinsic cellular properties. The same net-depressing pair rule, the same homeostatic budget and the same inhibition rescue almost every collapsing network built from one published conductance set and eliminate the majority of surviving networks built from another. Both signs are present inside one circuit model at one parameter setting.

The two signs do not share a cause. Within the seven conductance sets sampled here, elimination increased monotonically with intrinsic firing rate — but the same quantity **reverses** when the rate is moved inside a single set by injected current. **This is an ordering phenomenon across the sampled sets, not a rate law**: firing rate is a correlate across this sample and not a cause of elimination. Rescue does not track rate at all: driving a set until its control arm collapses as often as the follower’s does not make it rescuable, and substituting the follower’s conductances one at a time does not transfer the property. **Slow firing predicts fragility, but not rescue**.

The implications for the stabilising/destabilising literature are bounded by this model: **it is a simulation result about one circuit, and in that circuit the question does not have a single answer**. In this model a rule that is destabilising on one conductance set is protective on another that differs only in intrinsic conductances. Whether that carries to other models, other rules or living tissue is not addressed here.

The negative result of §2.6 is worth as much as the positive ones. It is common to expect that a network-level outcome traced to a cell’s intrinsic properties will localise to a channel, and here it does not at the resolution tested: the exhaustive single-substitution decomposition is empty at a resolution that detects the complete substitution at p = 2.3 × 10^−6^. We report this as an absence at a stated resolution, not as evidence that no such decomposition exists.

## 4. Limitations

1. **Everything here is computational**. No claim is made about living tissue. The preparation is a model culture built from published conductance sets; its wirings are realisations of one model, not independent biological preparations.
2. **One rule, one budget, one inhibition level for the main comparisons**. Initialisation and network size were tested and are reported in §6; **inhibition, the rule itself and the budget were not varied**, and nothing here establishes that the two signs survive changes to them.
3. **The relationship in §2.3 is carried by its low-rate end**. Excluding PY 1 it does not reach p < 0.05 at six cells, and the interval between 25.6 and 483 spikes/s contains no cell in the sampled catalogue.
4. **AB/PD 2’s elimination excess is not individually significant** (exact p = 0.125). It contributes a point estimate to §2.3, not a test.
5. **§2.3 is a correlation across a sample of seven, and §2.4 shows it reverses within a cell**. It supports no causal statement about firing rate.
6. **Four of the six single substitutions in §2.6 produced no measurable change in firing rate or bursting with the rule on**. Their nulls may reflect a preparation beyond the rule’s reach at that drive rather than a conductance that fails to confer rescue. We deliberately do not stratify the analysis on this: whether the rule “acted” is measured in the same arm as the rescue, and a rescued network fires more, so the split would be circular. It is a limit on the interpretation.
7. **The drives in §2.5 and §2.6 are not matched across arms**. I_H and I_leak required currents five to ten times further from baseline before their controls died, and the response of firing rate to injected current is not monotone in these substituted cells, so their qualifying drives were found by search rather than by ramp.
8. **The survival detector is a quorum on a fixed window**, and its horizon changes borderline classifications: two of 48 wirings reclassify between 30 s and 35 s. All results above are reported at 35 s, and on both cells at that horizon the registered and strict detectors agree on every wiring.
9. **The rank correlation treats each conductance set as one observation**. Per-cell denominators run from 3 to 36 wirings and are weighted equally; PY 1’s own elimination estimate is **0 of 3**, Wilson [0.000, 0.562]. Exact per-cell intervals are given in §2.3 and Figure 3, and the correlation is **not robust to precision weighting**, which we do not apply and do not claim.
10. **Multiple comparisons**. Eight arms were tested in §2.6 against a common baseline; the p-values reported are uncorrected, and a Bonferroni family-wise threshold at six informative arms is **0.0083**. The complete substitution (p < 10^−4^, and 2.3 × 10^−6^ pooled) clears it; no single substitution approaches it. The rate-matched pairwise comparisons in §2.6 were **chosen after seeing the control rates** and are reported as descriptive.
11. **The decomposition tests cardinality one and cardinality eight only**. Nothing between was run, so §2.6 establishes that no single conductance is sufficient and places an **upper bound of eight** on the size of a sufficient combination. It says nothing about two, three or seven.
12. **Each arm in §2.6 has n = 7 wirings**. The design is powered to detect a rescue against a zero baseline, not to estimate a small rescue rate.
13. **The phenomenon requires the network size reported here**. At 24 excitatory cells instead of 48, the follower-type population fires about four times faster and its control arm does not collapse, in either initialisation family, so there is no rescue to measure (§6). Everything in §2.2 and §2.5 is conditional on the larger network.
14. **The intrinsic conductances are fixed, and in real cultured neurons they are not**. Turrigiano, Abbott & Marder (1994) showed that cultured stomatogastric neurons change their intrinsic current densities in response to their own activity, on a timescale of days. Every result above treats a conductance vector as a constant property of a cell, so a real preparation would not necessarily stay at one point of §2.3 while the plasticity rule acted on it. This is a limitation of the model, not of the analysis, and it bears directly on the experiment proposed in §5.

## 5. A testable prediction, and what would retire the finding

The model predicts that in a preparation where population activity depends on recurrent excitation, a net-depressing plasticity rule under a conserved total conductance should have **opposite effects on survival in populations whose intrinsic firing rates differ by an order of magnitude at matched synaptic drive** — protective in the slow population, destructive in the fast one.

### We are not proposing that experiment yet

The programme’s own gate order puts a robustness test (§6) before any biological test, and that test has not run. What the model licenses today is the shape of the eventual experiment, recorded so that it is on the record before the enabling result rather than after: a preparation in which intrinsic excitability can be shifted pharmacologically or optogenetically within the same culture, with network survival scored under a fixed criterion and a plasticity manipulation applied in both states. Which preparation, and whether the model’s regime is reachable in it, is not settled here.

### The falsifier we would accept

in a preparation where the slow and fast conditions are produced within the same culture and the plasticity manipulation is identical, an effect of the same sign in both conditions retires the claim. So does the registered robustness test in §6 returning a sign change under a different initialisation or network size.

## 6. Robustness to initialisation and network size

The registered robustness test has now run. It asked whether the two-sign result of §2.2 survives changes of initialisation and network size: the same two conductance sets at matched fragility, across two initialisation families and two network sizes, at one inhibition level. Its falsifier, registered before the runs, was **a condition in which the two rescue rates cross**.

Family A is the initialisation used throughout this study — Gaussian jitter of 20% about the budget mean. Family B draws each initial weight uniformly on [0, 2·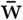], the same mean and roughly three times the spread, so the homeostatic budget starts identical.

| condition | follower-type rescue | pacemaker-type rescue | crossing |
| --- | --- | --- | --- |
| family A, 48 excitatory cells | 9/10 = 0.900 | 0/10 | **no**, $p = 1.2 \times 10^{-4}$ |
| family B, 48 excitatory cells | 9/9 = 1.000 | 0/10 | **no**, $p = 1.1 \times 10^{-5}$ |
| family A, 24 excitatory cells | — | 0/10 | **not testable** |
| family B, 24 excitatory cells | — | 0/10 | **not testable** |

**The falsifier did not fire in either condition where it could be tested**, and the initialisation family moves little: the control arm collapses in 10 of 11 and 9 of 11 wirings and the rule rescues 0.900 and 1.000.

**Two conditions could not be tested, and the reason is a limitation of the study rather than a null result**. At half the network size the follower-type population fires roughly four times faster — 105 and 100 spikes/s against 26 and 39 — and **its control arm does not collapse at all, in either initialisation family (0 of 11)**. With no collapse there is nothing to rescue and the conditional is undefined. The pacemaker-type set still collapses 10 of 10 at both sizes and is still rescued 0 of 10, so the loss is specific to the cell on which §2.2’s headline is measured.

**The collapse this study measures rescue against is therefore size-dependent**, and the result should be read as robust to initialisation at the size reported and as not occurring at half that size. That is a weaker statement than a sign reversal, and it is a different one: the phenomenon does not invert, it ceases to have an occasion.

## 7. Preregistration deviations, corrections, and audit trail

The protocol commits every prediction and falsifier before the corresponding run. More predictions failed than held; this section gives the rate and the failures that changed what the paper claims, and **the full table, prediction by prediction, is Supplementary Note S1**.

### 7.1 Predictions that failed

Nine of our registered predictions were refuted or proved unmeasurable, and three of the orchestrating record’s were refuted. Two of ours were registered *against* earlier predictions of ours after the evidence moved, and those two held. Three failures changed the manuscript rather than merely being recorded:

- **The mechanistic prediction was wrong twice**. We predicted that I_H, the rebound conductance, carried the rescue, on the argument that a rescue requires a cell able to fire from strong transient input; at a drive where its control collapsed it transferred nothing (0 of 7). Our second choice, I_KCa, reached 1 of 7. The orchestrating record’s prediction — a burst-duration conductance — also failed. §2.6 is what is left when all three mechanistic guesses fail.
- **A prediction about time that was simply lazy**. We predicted the pacemaker’s elimination fraction would rise at the longer horizon, reasoning that five more seconds is more time to collapse. It did not move at all (6 of 11 at both horizons): on that cell the control arm’s classification is horizon-invariant.
- **The horizon itself was a departure**. Two cells were first measured at a shorter horizon than the other five. That was found during revision, not registered in advance, and closed by re-measuring both at the common horizon with the consequence for this abstract of each outcome written before the run. The relationship in §2.3 was unchanged; the rescue conditional moved from 33/33 to 31/33, and the sentence it had supported — that not one network which collapsed without the rule collapsed with it — is no longer true and has been removed.

### 7.2 Design defects found and corrected, with what they cost

- **A statistic whose ceiling moves with the ordering variable**. (rescued − eliminated) cannot exceed the control’s death rate, which is what the ordering variable drives. Found after COLL-8 ran; the elimination fraction, which has no such ceiling, is reported as primary and orders the cells more strongly.
- **A circular covariate**. The per-wiring rate used to relate rate to death in COLL-9 was averaged over a window containing the death. The registered repair re-measures it over the deterministic first 5 s.
- **Two unreachable thresholds**. A falsifier set at “≥ 3 wirings dead in both arms” fires with probability 0.928 when the hypothesis it tests is true; its proposed replacement, a Wilson lower bound of 0.8, is unreachable at any feasible sample size (0.711 even at n = 100). Both were replaced with an exact test against p ≤ 0.5 before the run.
- **A minimum-denominator rule, registered in advance**, prevented an arm scoring 1 of 1 at p = 0.0455 from being reported as a finding. At a proper denominator that arm is 0 of 7. **A power calculation fed by the pilot that motivated it**. A sample-size estimate used a standard deviation from three seeds; at six seeds it fell by a third and the conclusion changed.
- **A rank correlation computed on ranks that did not handle ties**. Three of the seven cells in §2.3 tie at an elimination fraction of 0.000, and the rank function used broke those ties by input order — which, because the tied cells are the three slowest, is the tie-breaking most favourable to the hypothesis. On ranks with ties averaged, the standard definition, rho is **+0.852** rather than +0.893 and the six-cell value **+0.812** rather than +0.829. The quoted p = 0.0123 could not be reproduced under any tie convention or sidedness and has been replaced by the two-sided permutation p, 0.0286. **The inference is unchanged**: the seven-cell relationship is still significant at 0.05, the six-cell one still is not, and §2.3’s two stated limits stand. Found by STATS-1, which recomputes every statistic from the deposited runs; the figure and the figure checker were corrected with it.
- **Two p-values quoted from the one-sided tail while the rest were two-sided**. Found by recomputing every Fisher p in this manuscript from the deposited counts under both conventions. Only two of nine differed: §2.6’s I_KCa comparison (0.004 →0.005) and §6’s family A (6.0 × 10^−5^ →1.2 × 10^−4^). All are now two-sided, the convention is stated in Methods, and neither correction changes an inference.

### 7.3 A recorded discrepancy with the closest prior work

The work closest to ours is Jacquerie, Tyulmankov, Sacré & Drion, *Burst firing creates an attractor in synaptic weight dynamics* (PLOS Computational Biology 2026; our internal record cites it as 2025). For pair-based STDP in a feedforward network without homeostasis, they derive that bursting drives weights into a narrow attractor. In this preparation, measured at 50 ms resolution across five wirings and 60 s each, the coefficient of variation of recurrent excitatory weights **rises from 0.202 to 1.357** over 35 stored runs; it **grows inside every single burst — 214 of 214 — at +0.097 s**^**−1**^, and contracts only *between* bursts, where the homeostatic renormalisation acts, at **−0.013 s**^**−1**^, roughly seven times weaker, so the sign is inverted on both legs. **We did not observe the attractor-like convergence they describe**, under the present recurrent and homeostatically constrained conditions; **the tonic-firing control their result makes necessary was not run**, so this is a discrepancy we report and not a refutation we claim.

### Same substrate, opposite sign

We attribute the difference to the homeostatic budget and the recurrence, neither of which is in their derivation — but that attribution is an inference and not a measurement, because the tonic control arm their result makes obligatory **has not been run**. This discrepancy was recorded in our own literature review before this manuscript was drafted and is reported here because a failure list that omits it is not a failure list.

### 7.4 Results withdrawn

- A hysteresis signature reported on one ramp rate did not survive a slower ramp: the gap fell from 2.6× to 1.16× the noise and the feature moved one step along the parameter ladder, which identifies it as a lag rather than a coexistence. **No hysteresis claim is made in this manuscript**.
- An earlier generalisation from a single cell model to “self-generating networks” was withdrawn twice and is not used.

### 7.5 Execution record

Runs were interrupted three times by the machine rather than by the protocol: by repeated sleeps on battery, by the stopping rule, and by a low-memory event. **No data was lost in any of them**, because each lot is written to disk before its clock check. The execution environment was rebuilt once, mid-row, after a filesystem cleanup emptied it; a completed lot was then re-run and compared field by field against the stored version — **72 comparisons, 0 differences** — before any new numbers were accepted.

Per-run timings, the wall-versus-process-time divergences, the environment rebuild and its verification, and the batched-versus-unbatched acceptance gates are in **Supplementary Note S1** rather than here.

## 8. Methods

### 8.1 Shared with the companion manuscript

The neuron model, the simulated culture, the plasticity rule and the homeostatic budget are those of the companion preprint (Groppi & Ferrari, 2026; reference 17), and are summarised here only far enough to read the results. Each cell is the single-compartment STG model of Prinz, Billimoria & Marder (2003): eight Hodgkin–Huxley-type currents (I_Na, I_CaT, I_CaS, I_A, I_KCa, I_Kd, I_H, I_leak), an intracellular calcium pool (τ_Ca = 200 ms), and E_Ca recomputed each step from the Nernst equation with [Ca]_ext = 3 mM at 283 K. The gating kinetics are the voltage-clamp measurements of Turrigiano, LeMasson & Marder (1995) as formulated by Liu, Golowasch, Marder & Abbott (1998) and tabulated in Prinz et al. (2003, Table 1); the graded baseline synapses follow Prinz, Bucher & Marder (2004) after Abbott & Marder (1998). The dish is 48 excitatory and 12 inhibitory neurons, plastic excitatory→excitatory connections at probability 0.15, total incoming conductance budget fixed at (N−1)·p·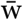with 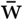, short-term depression (release fraction 0.20, recovery τ = 400 ms), and pair-based additive STDP with hard bounds w ∈ [0, 0.06], A_+_ = 0.012, τ_+_ = 16.8 ms, τ_−_ = 33.7 ms — the time constants measured by Bi & Poo (1998) — and A_−_ set from the area-ratio constraint so that the rule is net-depressing.

### 8.2 What is specific to this study

#### Validation of the preparation

The conductance sets are taken from a published database (Prinz, Bucher & Marder, 2004) and the circuit follows the STG modelling tradition (Abbott & Marder, 1998), but **this manuscript contains no independent validation that the simulated preparation reproduces a measured biological target**. Animal-to-animal variability in the real preparation is characterised by Bucher, Prinz & Marder (2005), and the degeneracy that motivates using a set of conductance vectors rather than one is the subject of Marder & Goaillard (2006). Neither comparison is made here, and the absence is a limitation of this study rather than of those sources.

#### Conductance sets

Excitatory populations are built from Prinz, Bucher & Marder (2004) Table 2 sets: AB/PD 1–5, LP 3 and PY 1. The inhibitory population is the LP 3 set throughout, including in the row where LP 3 is also the excitatory set; that row’s excitatory and inhibitory cells are therefore identical, which is stated in its registration and does not change its result.

#### Survival criterion and horizon

A network is alive if at least 24 of the 48 excitatory cells emit at least one spike in the final 10 s of the run. **Every result reported above is at a 35 s horizon**. Two cells were first measured at 30 s and re-measured at 35 s once the horizon was found to reclassify borderline wirings; the 30 s values survive here only in Limitation 8, which records that two of 48 wirings change classification between the horizons and that at 35 s the registered and strict detectors agree on every wiring of both cells. A stricter variant requiring the same quorum in the final 5 s alone is reported wherever the two differ.

#### Arms and wirings

Each cell contributes a rule-on and a rule-off arm on the *same* random wirings, so the 2 × 2 table is paired. Wirings are drawn from consecutive integer seeds and are not selected on any outcome.

#### Injected current

Where the drive is manipulated, a constant current is added to the excitatory population only, through the simulator’s existing external-current input; the inhibitory population is untouched. The current-to-rate relation is not monotone in the substituted cells of §2.6 and qualifying drives were found by search, with the search reported in the registration.

#### Validation of the simulation infrastructure

Runs are batched across wirings for speed, and the batched implementation is required to reproduce the unbatched one **bit for bit** — identical spike times and identical weights, not agreement to a tolerance — on both arms, at the cell and inhibition of the run in question. **This gate is re-run per cell rather than inherited, and passed at 0.000 × 10**^**0**^ **in every instance reported here**, including for the external-current parameter used in §2.4–§2.6, which no earlier row had exercised. Where a conductance substitution is expected to change nothing (the two channels identical in both cells) the substituted cell is likewise required to reproduce the unsubstituted one bit for bit, which is what establishes that the substitution machinery perturbs nothing by itself. These gates validate the implementation; they say nothing about whether the preparation matches biology, on which see above.

#### Statistics

Conditional probabilities carry Wilson 95% intervals, and every p-value quoted in the Results comes from the test named here for that quantity. Tests are exact throughout: binomial against p ≤ 0.5 for a single conditional, two-sided Fisher for 2 × 2 comparisons, McNemar’s exact form on discordant pairs where a sign is at issue, and full-enumeration permutation for Spearman correlations (5,040 permutations at seven cells). Spearman’s rho is computed on ranks with ties averaged, and its permutation p is two-sided, as the Fisher and McNemar p are. No asymptotic approximation is used, and no result rests on n < 3 in a denominator.

## 9. Pre-registration index

Every result traces to a section committed before its numbers existed, and the hashes below were verified against the repository when this manuscript was written. **Three departures from a clean ordering are listed under the table rather than smoothed over**, because an index that overstates itself is worth less than one that does not.

| result | § | registration | report |
| --- | --- | --- | --- |
| redistribution on driven cells | 2.1 | 234901f | f19c5f7 |
| elimination rate with an interval | 2.3 | 016f409 | be6b42e |
| rescue conditional, first measurement | 2.2 | 033aecb, gate d28c56b | 68ece31 |
| elimination with firing rate, five of the seven cells | 2.3 | 6d2a958 | 1bf9b3d |
| both endpoints at 35 s — **§2.2 and §2.3 report this row** | 2.2, 2.3 | 97f1254 | 74280de |
| robustness to initialisation and size | 6 | 8ac4db4 | 7e51296 |
| within-cell reversal | 2.4 | 6536bfd | d44e3f6 |
| fragility vs rescuability | 2.5 | 6536bfd, control cd835ce | d44e3f6 |
| decomposition, first half | 2.6 | 319f060 | ad1f380 |
| decomposition, second half | 2.6 | inside ad1f380 (see note i) | 59a7fc0 |
| decomposition, the two rate-raising channels | 2.6 | 60d8992 | 7ec5dd0 |

i. **The decomposition’s second half was registered inside a report commit**. `ad1f380` carries both the partial report of the first half and the registration of the second. Its registration content therefore precedes the second half’s numbers, which is what the protocol requires, but it does not sit in a commit of its own and a reader checking “registration before report” by commit order alone will see `ad1f380` and `59a7fc0` preceding `60d8992`.
ii. **One analysis script first appears in a report commit**. For the seven-cell ordering, `coll8b.py` and `analyse.py` are first committed in `1bf9b3d`, the report. The design, the grid and the thresholds were registered beforehand in `6d2a958`; the code was not.
iii. **The horizon change was not registered as such when it happened**. It followed from a detector limitation, and both endpoint cells were subsequently re-measured at the common horizon in a row whose registration (`97f1254`) states, before the run, what each possible outcome would do to this manuscript’s abstract. All figures above are at that single horizon.

Formatted in PLOS Computational Biology style. Every entry has been checked against the published record. Entries 1–17 are cited in the text; **entry 18 is listed only to disambiguate entry 16**, because the two share authors and year in our internal record and only one of them is the work §7.3 compares against.

## Author contributions

F.G. designed and ran the study, performed the analyses, and wrote the manuscript. J.F. contributed to the conception and interpretation of the work through continuous discussion and revised the manuscript. Both authors approved the final version.

## AI assistance statement

Generative AI tools (Claude Opus 5 for the executing sessions and Claude Fable 5.1 for the orchestrating session; Anthropic, 2026) were used under the authors’ direction for portions of code development, data-analysis support, figure-generation workflows, and manuscript drafting. In this study the AI system also executed the registered experimental protocol: it committed each registration, ran the simulations, and produced the reports from which this manuscript is written, under the design constraints and stopping rules recorded in the version history. The authors independently reviewed and verified the analyses, figures, claims, references, and final text and retain full responsibility for the work. No AI system is an author of this manuscript.

## Data and code availability

The simulator, the run scripts, the per-run result files, the figures with their source values and the registration record for this study are deposited at https://github.com/luviner-ai/luviner-neurolab, which also hosts the companion preprint’s code and data (Groppi & Ferrari, 2026; reference 17). The repository is archived on Zenodo under concept DOI **10.5281/zenodo.22287792**, which always resolves to the latest release; the release archiving the state of the repository at the time of this study is **v1.1.1, 10.5281/zenodo.22768648**, and the companion’s was v1.0.1, 10.5281/zenodo.22287793. Code is MIT-licensed; text, figures and data are CC BY 4.0.

Every reported result is generated by a committed analysis or simulation script and linked to the corresponding registration record in §9. The deposit includes `check_figures_paper2.py`, which re-derives every number appearing in a figure from the run outputs and compares it against this manuscript, and `check_stats_paper2.py`, which recomputes every count, interval and test quoted here from the same outputs and lists what it cannot reach. Both exit non-zero on any mismatch.

## Scope and boundary conditions

What this study establishes, and where it stops.

- **It establishes** that in this circuit model, at one inhibition and under one homeostatic budget, the same plasticity rule rescues most collapsing networks built from one published conductance set and eliminates the majority of surviving networks built from another.
- **It establishes** that within the seven sets sampled, elimination increases with intrinsic firing rate, and that this relationship reverses when the rate is moved inside a single set — so it is a correlate across the sample and not a cause.
- **It establishes** that slow firing predicts fragility and does not predict rescue, at matched control mortality.
- **It establishes** that no single conductance substitution confers the rescue, at a resolution that detects the complete substitution.
- **It does not establish** anything about living cultures. Every number is a model network, and the manuscript contains no validation of the preparation against a measured biological target.
- **It does not resolve** whether plasticity stabilises or destabilises recurrent circuits. It is a simulation result showing that, in this model, the question does not have one answer.
- **It does not identify a mechanism** for the rescue, and §2.6 is reported as an absence at a stated resolution.
- **It does not claim universality**. §2.1’s boundary case shows the effects depend on a preparation property that a given preparation may or may not have.
- **It is not a product claim**.

