## Supplementary Note S1 for "The same plasticity rule rescues networks built from one conductance set and destabilises networks built from another"

Audit tables and execution record for *The same plasticity rule rescues networks built from one conductance set and destabilises networks built from another*. Everything here is referenced from §7 of the main text and is separated from it because it is a record, not an argument.

### S1.1 Every registered prediction that was scored

#### 7.1 Predictions that failed — ours

| prediction | row | outcome |
|---|---|---|
| No sign flip in (rescued − eliminated) across inhibition | COLL-5 | **held**; the registered magnitude at g_ie 0.045 (−0.25 to −0.33) was wrong, measured −0.417 |
| Rescue conditional on opportunity lands in [0.45, 0.70] | COLL-6 | **refuted** — 1.000 |
| Rescue is lower among the earliest-dying wirings | COLL-6 | **unmeasurable** — 32 of 33 wirings share one value of the covariate; the split had no variance |
| Elimination fraction on healthy wirings in [0.2, 0.5] | COLL-6 | **unmeasurable** — denominator of 3 |
| Four of five new cells are protective | COLL-8 | **refuted on all five** |
| Rescue follows wherever the opportunity exists | COLL-9 | **refuted** — 0 of 21 |
| I_H carries the rescue | COLL-11 | **refuted** — 0 of 7 at a qualifying drive |
| I_KCa carries the rescue | COLL-11 | **not established** — 1 of 7, p = 0.25 |
| No single conductance carries the rescue | COLL-11 | **held** — registered against our own I_H prediction after the first half's numbers arrived |
| Rescue lands at 31/33 at the longer horizon | COLL-14 | **held, exactly** |
| The three control-surviving wirings stay alive | COLL-14 | **held** — 0 of 3 |
| The pacemaker's elimination rises at the longer horizon | COLL-14 | **refuted** — 6/11 unchanged; the reasoning ("more time means more deaths") was wrong, the control arm's classification is horizon-invariant on that cell |
| The relationship in §2.3 survives at one horizon | COLL-14 | **held** — +0.852, p = 0.0286, identical (the value corrected in §7.2; the relationship is unchanged by the horizon either way) |

#### 7.2 Predictions that failed — the orchestrating record's

| prediction | row | outcome |
|---|---|---|
| The sign flips with inhibition, protective below the band | COLL-5 | **refuted** — the quantity goes from 0 to negative, not from positive to negative |
| Cell identity orders the effect: LP and PY followers protective | COLL-8 | **refuted on LP 3**, the cell named in advance to decide it |
| A burst-duration conductance carries the rescue | COLL-11 | **refuted** — I_CaS 0/7, I_KCa 1/7 |

### S1.2 Execution record, run by run

Runs were interrupted three times by the machine rather than by the protocol: by repeated sleeps on battery power, by a stopping rule, and by a low-memory event. No data was lost in any of them, because each lot is written to disk before its clock check.

One row executed across repeated machine sleeps: 938 minutes of wall clock for 43.7 minutes of process time. The in-script clock check detected the divergence from the operating system's sleep log, and the stopping rule, which is written on process time rather than wall time, was unaffected.

The execution environment was rebuilt once, mid-row, after a filesystem cleanup emptied it — the package directories survived while their Python sources were deleted, so an existence check passed and imports failed. The environment was rebuilt at identical interpreter and package versions, and a completed lot was then deleted, re-run under the new environment and compared field by field against the stored copy: **12 wirings x 6 fields = 72 comparisons, 0 differences**, identical spike for spike. Only then were new numbers accepted.

Two rows stopped on their stopping rule with one condition outstanding and were completed in a continuation; both are noted where they occur in the main text.

### S1.3 Batched-versus-unbatched acceptance gates

Every cell and every parameter regime used in the main text was gated by requiring the batched implementation to reproduce the unbatched one bit for bit, on spikes and on weights, on both arms. Every such gate passed at 0.000 x 10^0. The gate was re-run per cell rather than inherited, and it was extended to the external-current parameter when that parameter was first used, and to the conductance-substitution machinery by requiring that substituting a channel identical in both cells changes nothing.

### S1.4 Every wiring at both horizons

The two cells of §2.2 were first measured at 30 s and re-measured at 35 s. This is the comparison wiring by wiring; the manuscript reports the 35 s column throughout. Only wirings whose classification changed are listed individually — the rest are identical on both arms at both horizons and are summarised by count.

**PY 1**, 36 wirings, seeds 30–65.

| | **30 s** | **35 s** |
|---|---|---|
| control collapsed | 33/36 | 33/36 |
| rescued given the control collapsed | 33/33 | 31/33 |
| eliminated given the control survived | 0/3 | 0/3 |
| wirings unchanged on both arms | — | **34 of 36** |

Wirings that changed classification:

| **seed** | **control 30 s** | **rule 30 s** | **control 35 s** | **rule 35 s** |
|---|---|---|---|---|
| 32 | collapsed | alive | collapsed | collapsed |
| 57 | collapsed | alive | collapsed | collapsed |

**AB/PD 2**, 12 wirings, seeds 30–41.

| | **30 s** | **35 s** |
|---|---|---|
| control collapsed | 1/12 | 1/12 |
| rescued given the control collapsed | 1/1 | 1/1 |
| eliminated given the control survived | 6/11 | 6/11 |
| wirings unchanged on both arms | — | **12 of 12** |

No wiring changed classification on either arm.

Both detectors were recorded for every wiring at 35 s — the registered one (any spike from the quorum in the final 10 s) and a stricter variant (the quorum in the final 5 s alone). **They agree on every wiring of both cells at that horizon**, which is why the main text reports a single classification. The two wirings that changed above are the two that already failed the strict detector at 30 s, and they were named as the candidates in the registration of the re-measurement before it ran.
